# De novo assembly of viral quasispecies using overlap graphs

**DOI:** 10.1101/080341

**Authors:** Jasmijn A. Baaijens, Amal Zine El Aabidine, Eric Rivals, Alexander Schönhuth

**Author notes:** These authors contributed equally (Eric Rivals, Alexander Schönhuth).

## Abstract

A viral quasispecies, the ensemble of viral strains populating an infected person, can be highly diverse. For optimal assessment of virulence, pathogenesis and therapy selection, determining the haplotypes of the individual strains can play a key role. As many viruses are subject to high mutation and recombination rates, high-quality reference genomes are often not available at the time of a new disease outbreak. We present SAVAGE, a computational tool for reconstructing individual haplotypes of intrahost virus strains without the need for a high-quality reference genome. SAVAGE makes use of either FM-index based data structures or ad-hoc consensus reference sequence for constructing overlap graphs from patient sample data. In this overlap graph, nodes represent reads and/or contigs, while edges reflect that two reads/contigs, based on sound statistical considerations, represent identical haplotypic sequence. Following an iterative scheme, a new overlap assembly algorithm that is based on the enumeration of statistically well-calibrated groups of reads/contigs then efficiently reconstructs the individual haplotypes from this overlap graph. In benchmark experiments on simulated and on real deep coverage data, SAVAGE drastically outperforms generic *de novo* assemblers as well as the only specialized *de novo* viral quasispecies assembler available so far. When run on ad-hoc consensus reference sequence, SAVAGE performs very favorably in comparison with state-of-the-art reference genome guided tools. We also apply SAVAGE on two deep coverage samples of patients infected by the Zika and the hepatitis C virus, respectively, which sheds light on the genetic structures of the respective viral quasispecies.

## Introduction

Viruses such as HIV, the Zika and the Ebola virus, populate their hosts as an ensemble of genetically related but different mutant strains, commonly referred to as *viral quasispecies*. These strains, each characterized by its own haplotypic sequence, are subject to high mutation and recombination rates (Domingo et al. 2012; Duffy et al. 2008). Sequencing methods aim at capturing the genetic diversity of viral quasispecies present in infected samples; the promise is that next-generation sequencing (NGS) based methods will assist clinicians in selecting treatment options and other clinically relevant decisions.

Ideally, a *viral quasispecies assembly* characterizes the genetic diversity of an infection by presenting all of the viral haplotypes, together with their abundance rates. There are two major challenges in this.

(1) The number of different strains is usually unknown. Furthermore, two different strains can differ by only minor amounts of distinguishing mutations. Last but not least, abundance rates can be as low as the sequencing error rates, which hampers the detection of true mutations present at low frequency.

(2) Due to the great diversity and the high mutation rates, reference genomes representing high-quality consensus genome sequences can be obsolete at the time of the disease outbreak. The lack of a suitable reference genome is a major hindrance for many viral quasispecies assembly approaches.

It is important to understand that all existing assembly methods fail to address either the first or the second point. Recent *reference-guided approaches* specialized in viral quasispecies assembly suggested statistical frameworks modeling the driving forces underlying the evolution of viral quasispecies. While previous approaches focused mostly on local reconstruction of haplotypes (Huang et al. 2012; Quince et al. 2011; Zagordi et al. 2011; 2010), more advanced approaches aimed at global reconstruction of haplotypes, for example, by making use of Dirichlet process mixture models (Prabhakaran et al. 2014), hidden Markov models (Töpfer et al. 2013), or sampling schemes (Prosperi and Salemi 2012). There are also recent combinatorial approaches which compute paths in overlap graphs (Astrovskaya et al. 2011), enumerate maximal cliques in overlap graphs (Töpfer et al. 2014), or compute maximal independent sets in conflict graphs (Mangul et al. 2014). While these approaches soundly address point (1), the vast majority of them depends on high-quality reference sequence as a backbone to their methods, which in turn is the reason why they fail to address (2). Hence, when confronted with hitherto unknown, significantly deviating mutation patterns, these approaches fail to perform sufficiently well.

On the other hand, *de novo assembly approaches* do not depend on reference genomes. Although there exist numerous *de novo* approaches for mammalian genome assembly, – see e.g. (Bradnam et al. 2013; Gurevich et al. 2013; Salzberg et al. 2011) for comparative evaluations – these generic methods are not well suited for the viral quasispecies assembly problem. The key difference is that mutation rates in viruses are orders of magnitude higher than in eukaryotes, resulting in multiple polymorphic sites within a single read (Domingo et al. 2012; Duffy et al. 2008). This makes it possible to phase mutations into separate haplotypes; however, generic assembly approaches do not exploit this property. Rather, generic assemblers aim at reconstructing one single consensus sequence or are not designed to handle genomes of heavily polyploid organisms. In this regard, note that there are *de novo* assemblers that specialize in viral genome assembly already (Yang et al. 2012; Hunt et al. 2015). However, also these specialized approaches aim at assembling consensus genomes rather than strain-specific sequence, where the goal is to construct new reference rather than individual sequence. To our knowledge, the only existing *de novo* approach for haplotype-resolved viral quasispecies assembly is MLEHaplo (Malhotra et al. 2016b). However, as our evaluations will demonstrate, MLEHaplo does not even compare favorably with generic de novo assemblers. As a consequence, while addressing (2), existing de novo assembly methods fail to address point (1) to a satisfactory degree.

A possible principled issue is that nearly all of the NGS based genome assemblers, including the abovementioned specialized *de novo* viral quasispecies approaches, rely on the *de Bruijn graph* as assembly paradigm. Thereby, reads are decomposed into *k*-mers, where *k* is usually considerably smaller than the read length. As a generalization of this concept the paired de Bruijn graph has been introduced (Medvedev et al. 2011), which incorporates mate pair information into the graph structure itself instead of analyzing mate pairs in a post-processing step, which yields larger contigs in the assembly. As mentioned above, it is imperative in viral quasispecies assembly to distinguish low-frequency mutations from sequencing errors. While low-frequency mutations are genetically linked, hence co-occur within different reads, sequencing errors do not exhibit patterns of co-occurrence. The detection of patterns of co-occurrence is decisively supported by examining reads at their full length, but this information cannot be exploited with de Bruijn graphs. Overlap graphs on the other hand make use of full-length reads and do not decompose them into smaller parts; hence, we reason that the overlap graph paradigm suits the problem of viral quasispecies assembly better.

The only existing method for viral quasispecies assembly based on overlap graphs is HaploClique (Töpfer et al. 2014). Although this method is reference guided, it uses the reference solely for providing anchor points for constructing an overlap graph. Unlike in many other approaches (Di Giallonardo et al. 2014; Töpfer et al. 2013; Zagordi et al. 2010; 2011), the haplotype sequences are then assembled from the reads, and not from the reference. While providing inspiration in general, the HaploClique algorithm has proven too expensive—already data sets of about 1000x coverage require excessive computational resources. The reason is that it is based on the enumeration of maximal cliques, which is exponential in the read coverage, both in terms of runtime and space. We therefore present a novel, more efficient algorithm for the clique enumeration part of the assembly algorithm.

There are two exit strategies to resolve the issue of the possible lack of a reference genome. The *first strategy* is to construct consensus genome sequence from the patient samples themselves, using one of the available *de novo* consensus genome assemblers (among which, the most popular tool is VICUNA (Yang et al. 2012)), and to subsequently run one of the reference-guided approaches using this ad-hoc consensus as a reference. This strategy has also been suggested by (Mangul et al. 2014) and we shall further explore it here. The *second strategy* is to construct an overlap graph directly from the patient sample reads. Subsequently, one employs a ploidy-aware assembly algorithm that can extract strain-specific sequences from overlap graphs. The challenge is that constructing overlap graphs requires a pairwise comparison of all reads, which, for deep coverage data sets, requires sophisticated indexing techniques to be feasible. Here, we show how to make efficient use of FM-index based techniques (Välimäki et al. 2012) to construct overlap graphs without any need for a reference genome. As such, we provide *the first approach for de novo assembly of viral quasispecies based on overlap graphs*.

In summary, we make relevant contributions for

i. the construction of overlap graphs from deep coverage read data and
ii. viral quasispecies assembly using the overlap graph assembly paradigm.

In combination, we present SAVAGE (Strain Aware VirAl GEnome assembly), a method that allows for reference-free assembly of viral quasispecies from sequencing data sets of deep coverage (20 000x and more). In this, we do not only provide the first genuine *de novo* viral quasispecies assembly approach based on overlap graphs, but we also provide the first method that can exploit ad-hoc consensus sequence generated from patient samples, as computed for example by VICUNA (Yang et al. 2012), for high-performance viral quasispecies assembly.

## Results

We have designed and implemented SAVAGE (Strain Aware VirAl GEnome assembly), a method for *de novo* viral quasispecies assembly based on overlap graphs. In this section, we provide a high-level description of the algorithmic approach and analyze its performance, also in comparison to state-of-the-art viral quasispecies assembly tools and several established generic genome assemblers. Finally, we present assembly results using SAVAGE on two real virus samples from patients infected by the Zika virus and hepatitis C virus, respectively. We refer to the Methods section for any methodological details.

### Approach

Our algorithm proceeds in three stages (panel A of Figure 1), each of which iteratively clusters the input sequences and extends them to unique haplotypes. While *Stage a* has the original reads as input and contigs as output, *Stage b* has these contigs as input and maximally extended contigs as output. The extended contigs are supposed to reflect individual haplotype sequences. Finally, the optional *Stage c* merges maximally extended contigs into master contigs, each representing a group of very closely related strains. This reflects the existence of **master strains** in many viruses, where each individual haplotype deviates from one of the master strains by only a relatively minor amount of mutations (the ensemble of which is commonly referred to as *mutant class* in the literature and reflects a viral subpopulation, see e.g. Domingo et al. (2012)). Each stage is divided into **overlap graph construction** (upper part of panel C in Figure 1) and **overlap graph based assembly** (lower part of panel C in Figure 1). Between the stages, this generic structure only differs in the details.

**Figure 1:**
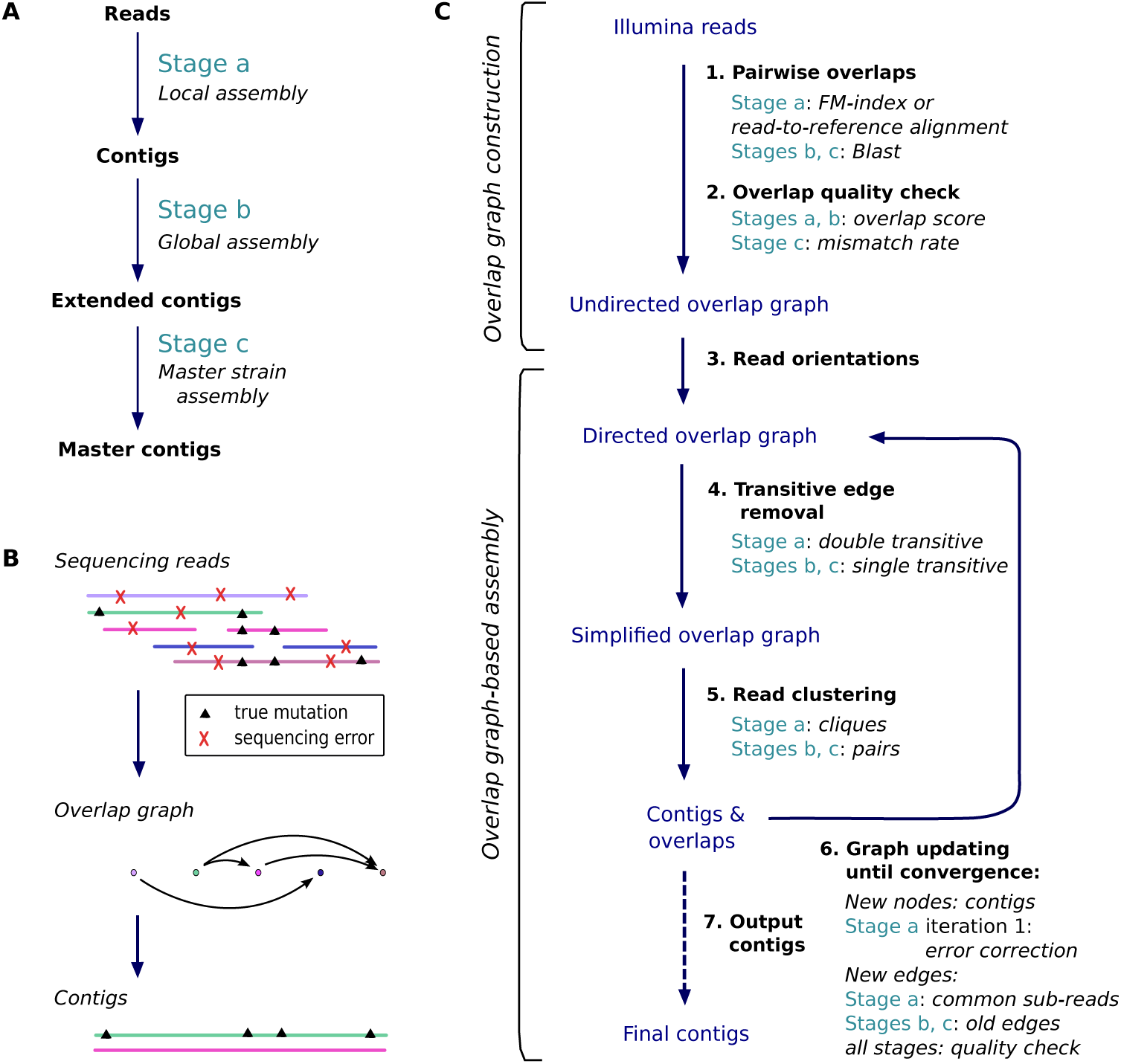
An overview of the workflow and algorithms of SAVAGE. **A.** The three stages of SAVAGE. Each assembles sequences into longer sequences. For clarity, we assign different names to the sequences output by each stage: contigs, maximally extended contigs, and master contigs, respectively. **B.** Principle of overlap graph construction and distinction among the reads between errors and shared mutations. **C.** Each stage has two steps: first, the overlap graph construction, second, assembly. This panel summarizes the differences in each step between the three stages. During overlap graph-based assembly, steps 4 to 6 are repeated iteratively until there are no edges left in the overlap graph.

The strength of overlap graphs for viral quasispecies assembly is in identifying co-occurring mutations, thus enabling the phasing of mutations from the same strain. We distinguish sequencing errors from true mutations by posing very strong constraints on the overlaps in terms of minimal overlap length and sequence similarity. In addition, we make use of paired-end read information. This results in a very conservative overlap graph, where an edge indicates that two sequences are very likely to originate from the same virus strain. Therefore, by enumerating cliques in the overlap graph we cluster the reads per strain, thus reconstructing the individual haplotypes of the viral quasispecies.

We construct overlap graphs in two steps: first, pairs of reads are determined that share sufficiently long and well-matching overlaps, followed by a statistical evaluation of the quality of each overlap. We explore two options for finding all such overlap candidates. The first option is to apply a completely *de novo* procedure using FM-index based techniques (Välimäki et al. 2012). The second option is to align all reads against a reference genome, such that read-to-read alignments can be induced from the read-to-reference alignments. However, in case of a viral outbreak there may not be a suitable reference genome available; we target such cases by constructing an ad-hoc consensus sequence from the patient samples, as computed by VICUNA (Yang et al. 2012).

SAVAGE offers three different modes, corresponding to the different approaches to overlap graph construction described above: **SAVAGE-de-novo** uses the first option and is therefore completely reference-free, while **SAVAGE-b-ref** uses the second option and thus relies on a bootstrap reference sequence. For benchmarking purposes we also consider **SAVAGE-h-ref**, which takes as input an existing, high quality reference sequence.

### Benchmarking data

For benchmarking experiments and performance analysis, we considered several simulated data sets, one gold standard benchmark from real sequencing reads, and two real patient samples. For the simulated data sets, sequencing reads were created using the simulation software SimSeq (see Methods).

#### Simulated benchmarks

We created five simulated data sets for benchmarking, consisting of 2×250bp Illumina MiSeq reads and representing quasispecies infections from different viruses: human immunodeficiency virus (HIV), hepatitis C virus (HCV), and Zika virus (ZIKV). We varied the number of strains per sample, as well as the relative abundances of those strains and the pairwise divergence between strains. To get data sets as realistic as possible, we used true viral genomes from the NCBI database and Illumina MiSeq error profiles during simulations. Characteristics of each benchmark are given in Table 1 and additional information can be found in Supplemental Methods.

**Table 1:** Characteristics of benchmarking data sets. For each benchmark we specify virus type, genome length, average coverage, strain count, relative abundance, and pairwise divergence. For the 600x HIV mix, the strains were homogeneously distributed with a relative abundance of 20% each.

| Data set | Virus type | Genome length (bp) | Average coverage | Strain count | Strain abundance | Pairwise divergence |
| --- | --- | --- | --- | --- | --- | --- |
| 600x HIV mix | HIV-1 | 9478 – 9719 | 600x | 5 | 20 % | 1 – 6 % |
| 5-strain HIV mix | HIV-1 | 9478 – 9719 | 20 000x | 5 | 5 – 28 % | 1 – 6 % |
| 10-strain HCV mix | HCV-1a | 9273 – 9311 | 20 000x | 10 | 5 – 19 % | 6 – 9 % |
| 3-strain ZIKV mix | ZIKV | 10251 – 10269 | 20 000x | 3 | 16 – 60 % | 3 – 10 % |
| 15-strain ZIKV mix | ZIKV | 10251 – 10269 | 20 000x | 15 | 2 – 13 % | 1 – 10 % |
| *Lab mix* | HIV-1 | 9478 – 9719 | 20 000x | 5 | 10 – 30 % | 1 – 6 % |

#### Lab mix

In addition to the simulated benchmarks, we also considered a real Illumina MiSeq (2×250 bp) data set with an average coverage of ∼20 000x, obtained from a lab mixture of five HIV strains (see also Table 1). This data set was recently presented as a gold standard benchmark (Di Giallonardo et al. 2014); we will refer to it as the *lab mix*.

#### Divergence-vs-ratio

To analyze the combined effect of the levels of divergence and of the relative abundance of the strains, we constructed 36 additional data sets as follows. Starting from the HIV-1 89.6 haplotype, we created six alternative haplotypes by introducing, respectively, 0.5%, 0.75%, 1%, 2.5%, 5%, and 10% random mutations. For each of those six alternative strains, we created six data sets by simulating reads (2×250 bp Illumina MiSeq) from the mutated strain and the original at a ratio of 1:1, 1:2, 1:5, 1:10, 1:50, and 1:100, respectively, with a total coverage of 500x per data set.

#### Zika virus sample

We applied SAVAGE to a sample of Asian-lineage Zika virus (ZIKV) consisting of Illumina MiSeq 2×300 bp sequencing reads (∼30,000× coverage) obtained from a rhesus macaque after four days of infection (Dudley et al. 2016, animal 393422).

#### Hepatitis C virus sample

In addition to the Zika virus sample, we also used a hepatitis C virus (HCV) sample of approximately 80 000x coverage, covering a region of ∼3000bp containing the HS5B gene.

### Evaluation preliminaries

In the case of a viral outbreak, the agent and its genome may be unknown (or may have significantly diverged from closely related strains such that available reference sequences are potentially inadequate for analysis), and the samples taken from infected patients contain an unknown number of divergent strains. Here, we target these cases where no reference genome is available. A sample sequenced with Next Generation Sequencing technology delivers enough reads and sufficient coverage to allow a *de novo* assembly of a viral genome (here, we mean a single genome assembly, not a quasispecies assembly), to be used as an ad-hoc reference genome for further analyses. However, such genome sequences may not represent any of the true viral haplotypes present in the sample sufficiently well.

In the remainder of this paper, all assembly algorithms were run using default settings. Evaluations of assemblies were performed with MetaQUAST (Mikheenko et al. 2016), which computes the usual statistics – number of contigs, largest contigs, N50, misassembled contig length, target genome(s) covered, and error rates – and we accounted only for contigs larger than a threshold of 500 bp. A contig is called misassembled if it contains at least one misassembly, i.e., a position where the left and right flanking sequences align to the true genomes with a gap or overlap of more than 1 kbp, or align to different strands, or even align to different strains.

We compare *de novo* methods and reference-guided approaches. While *de novo* algorithms proceed by iteratively extending contigs until some convergence criterion is met, reference-guided approaches alter the reference sequence until a set of haplotypes is obtained that is supposed to represent the quasispecies. By altering the reference genome, all output sequences have the same length, which means that the N50 score equals the length of the output sequences. For *de novo* approaches, on the other hand, the N50 score provides an indication of the contig length distribution.

### Failure of existing de novo assemblers on low-frequency strains

We explored the ability of generic genome assemblers to reconstruct a viral quasispecies. From the broad collection of tools available, we selected four assemblers: SGA (Simpson and Durbin 2012), SOAPdenovo2 (Luo et al. 2012), SPAdes (Bankevich et al. 2012), and metaSPAdes (Nurk et al. 2016). The first two methods, SGA and SOAPdenovo2, are generic assemblers, mostly used on mammalian genomes. SPAdes was originally designed for bacterial genomes, and metaSPAdes is a version of SPAdes adapted for metagenome assembly.

First, we evaluate performance on all simulated benchmarks. Table 2 presents results for all methods on the 5-strain HIV mix, the 10-strain HCV mix, and the 15-strain ZIKV mix. The only method capable of assembling at least half a viral quasispecies on a 20 000x simulated data set is SPAdes, the only close alternative being metaSPAdes with 45.9% on the 10-strain HCV mix. For the 5-strain HIV mix and the 10-strain HCV mix, SPAdes assembles 91.3–91.7% (SAVAGE-de-novo: ≥ 99.6%) of the true viral genomes at an error rate of 0.015 – 0.084% (SAVAGE-de-novo: 0.004%), showing that SPAdes misses to assemble a considerable fraction of the quasispecies. This becomes more evident on the 15-strain ZIKV mix, which contains several low-frequency strains: SPAdes only recovers 65.6% of the target genomes (SAVAGE-denovo: 99.4%). The explanation for this is that SPAdes misses to assemble strains of low frequency, as Figure 2 further reveals: here, a comparison of all approaches is shown when at most a bootstrap reference is provided. The performance of each approach is evaluated on each of the strains of the 20 000x benchmarks from Table 1 individually, and results are stratified by the relative abundances of the strains. We see that SPAdes recovers only 46.8% of the strains of frequency of less than 5%.

**Table 2:** Assembly results per method on simulated HIV, HCV, and ZIKV benchmarks (20 000x coverage). For reference guided methods we present results using an established, high quality reference genome (h-ref) as well as an ad-hoc, bootstrap reference genome (b-ref). All assemblies were evaluated on the following criteria: number of contigs ≥ 500 bp, length of the largest contig, N50 statistic, MissAssembled Contigs (MAC) length relative to total contig length, percentage of the target genomes recovered, percentage of undetermined bases (N), and percentage of mismatches and indels compared to the ground truth.

|  | # contigs<br>≥ 500bp | largest<br>contig | N50 | MAC<br>length<br>(%) | target<br>genomes<br>(%) | N-rate<br>(%) | mismatches<br>(%) | indels<br>(%) |
| --- | --- | --- | --- | --- | --- | --- | --- | --- |
| **5-strain HIV mix** |  |  |  |  |  |  |  |  |
| PredictHaplo-h-ref | 5 | 9720 | 9720 | 0 | 99.6 | 0.603 | 0.085 | 0.102 |
| PredictHaplo-b-ref | 5 | 9578 | 9578 | 0 | 93.8 | 0.284 | 0.110 | 0.104 |
| ShoRAH-h-ref | 289 | 9514 | 9514 | 0 | 39.4 | 0.268 | 2.403 | 0.016 |
| ShoRAH-b-ref | 242 | 9501 | 9501 | 7.0 | 93.8 | 0.127 | 3.197 | 0.124 |
| SAVAGE-de-novo | 36 | 9413 | 4913 | 0 | 99.8 | 0 | 0.004 | 0 |
| SAVAGE-h-ref | 28 | 9634 | 5027 | 0 | 99.6 | 0 | 0.004 | 0 |
| SAVAGE-b-ref | 59 | 9463 | 2424 | 0 | 99.5 | 0.002 | 0.071 | 0.002 |
| SGA | 36 | 1034 | 650 | 0 | 32.4 | 0 | 1.294 | 0.026 |
| SOAPdenovo2 | 36 | 844 | 516 | 0 | 35.7 | 0 | 0.633 | 0 |
| SPAdes | 14 | 9789 | 5873 | 0 | 91.7 | 0 | 0.084 | 0.002 |
| metaSPAdes | 13 | 7044 | 5159 | 0 | 32.7 | 0 | 1.681 | 0.013 |
| **10-strain HCV mix** |  |  |  |  |  |  |  |  |
| PredictHaplo-h-ref | 9 | 9313 | 9313 | 0 | 90.0 | 0.004 | 0.402 | 0.010 |
| PredictHaplo-b-ref | 9 | 7636 | 7636 | 0 | 73.8 | 0.006 | 0.053 | 0 |
| ShoRAH-h-ref | - | - | - | - | - | - | - | - |
| ShoRAH-b-ref | 639 | 7570 | 7570 | 0 | 56.9 | 0 | 4.381 | 0.011 |
| SAVAGE-de-novo | 46 | 9297 | 8248 | 0 | 99.6 | 0.002 | 0.004 | 0 |
| SAVAGE-h-ref | 85 | 9247 | 3716 | 0 | 99.6 | 0 | 0.004 | 0 |
| SAVAGE-b-ref | 84 | 7802 | 2943 | 0 | 86.0 | 0 | 0.001 | 0 |
| SGA | 33 | 832 | 638 | 0 | 18.1 | 0 | 1.439 | 0 |
| SOAPdenovo2 | 41 | 926 | 531 | 0 | 22.0 | 0 | 0.551 | 0 |
| SPAdes | 13 | 9311 | 8582 | 0 | 91.3 | 0 | 0.015 | 0 |
| metaSPAdes | 81 | 3041 | 1549 | 0 | 45.9 | 0 | 2.133 | 0 |
| **15-strain ZIKV mix** |  |  |  |  |  |  |  |  |
| PredictHaplo-h-ref | 8 | 10258 | 10258 | 0 | 53.3 | 0.032 | 0.147 | 0.046 |
| PredictHaplo-b-ref | 8 | 10270 | 10270 | 0 | 53.3 | 0.001 | 0.121 | 0.004 |
| ShoRAH-h-ref | - | - | - | - | - | - | - | - |
| ShoRAH-b-ref | 493 | 10117 | 10117 | 0 | 26.3 | 0.053 | 4.403 | 0.017 |
| SAVAGE-de-novo | 607 | 9282 | 2103 | 0 | 99.4 | 0.002 | 0.016 | 0 |
| SAVAGE-h-ref | 641 | 10243 | 1935 | 0 | 99.4 | 0.002 | 0.006 | 0 |
| SAVAGE-b-ref | 604 | 9079 | 2018 | 0 | 99.5 | 0.002 | 0.011 | 0 |
| SGA | 0 | - | - | - | 0 | - | - | - |
| SOAPdenovo2 | 56 | 1025 | 562 | 0 | 21.0 | 0 | 0.545 | 0 |
| SPAdes | 60 | 10269 | 2577 | 0 | 65.6 | 0 | 0.131 | 0 |
| metaSPAdes | 37 | 6495 | 3926 | 0 | 17.5 | 0 | 1.200 | 0 |

**Figure 2:**
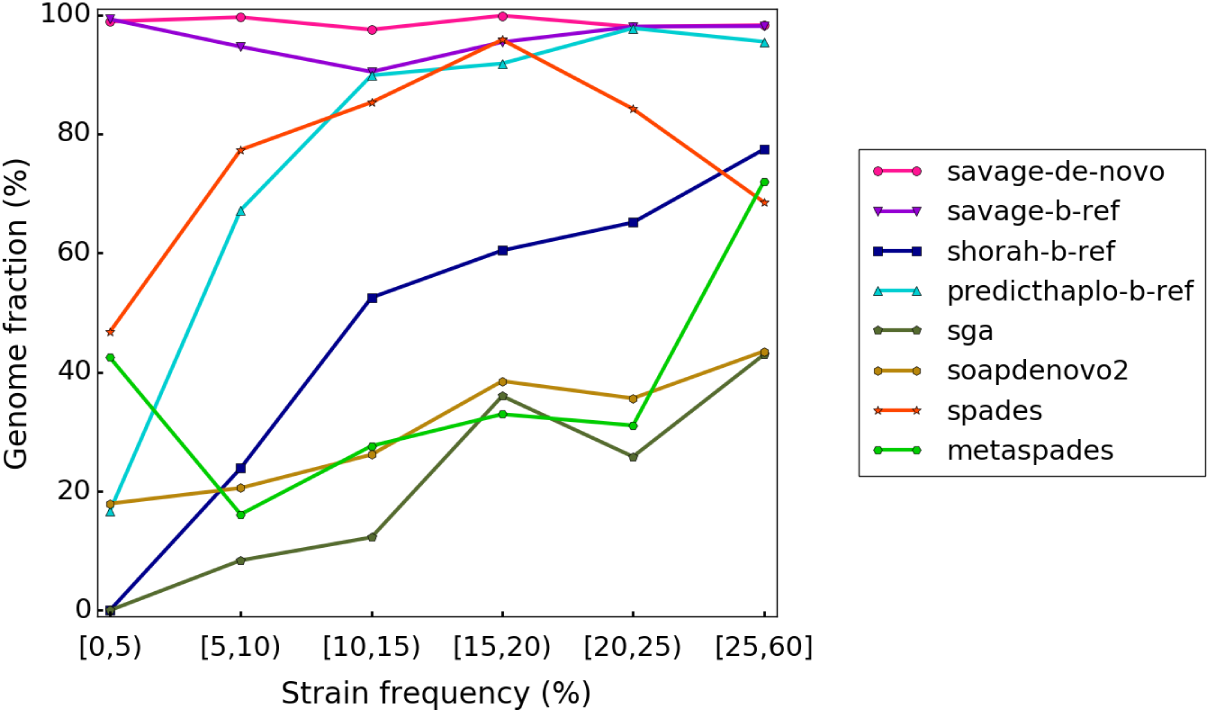
Target genome fraction recovered per strain for all 20 000x benchmarks, stratified by strain frequency.

Similar results for the 600x HIV mix and the 3-strain ZIKV mix can be found in Supplemental Tables S1 and S2; these are relatively easy data sets, since neither contains any low-frequency strains. Both SOAPdenovo2 and SPAdes perform reasonably on the 600x data set, reconstructing 78.9% and 87.8% of the viral quasispecies, respectively. SGA and metaSPAdes, on the other hand, do not recover more than 19% of the quasispecies. For the 3-strain ZIKV mix, only SPAdes is able to reconstruct more than 40% of the quasispecies; in fact, it finds 99.6% of the target genomes, performing almost perfectly on this low-ploidy dataset, which is no surprise because assemblers like SPAdes generally target at genomes of limited ploidy.

Finally, we consider the lab mix, which is based on real data and hence the most challenging benchmark. Table 3 presents results for all methods. SGA, SOAPdenovo2, SPAdes, and metaSPAdes all perform quite similarly, reconstructing only 41.0–53.7% of the viral quasispecies at very high error rates (1.1–2.0%). This shows that each of these assemblers has difficulty distinguishing sequencing errors from true variants, thus pointing out the need for specialized viral quasispecies assemblers.

**Table 3:**
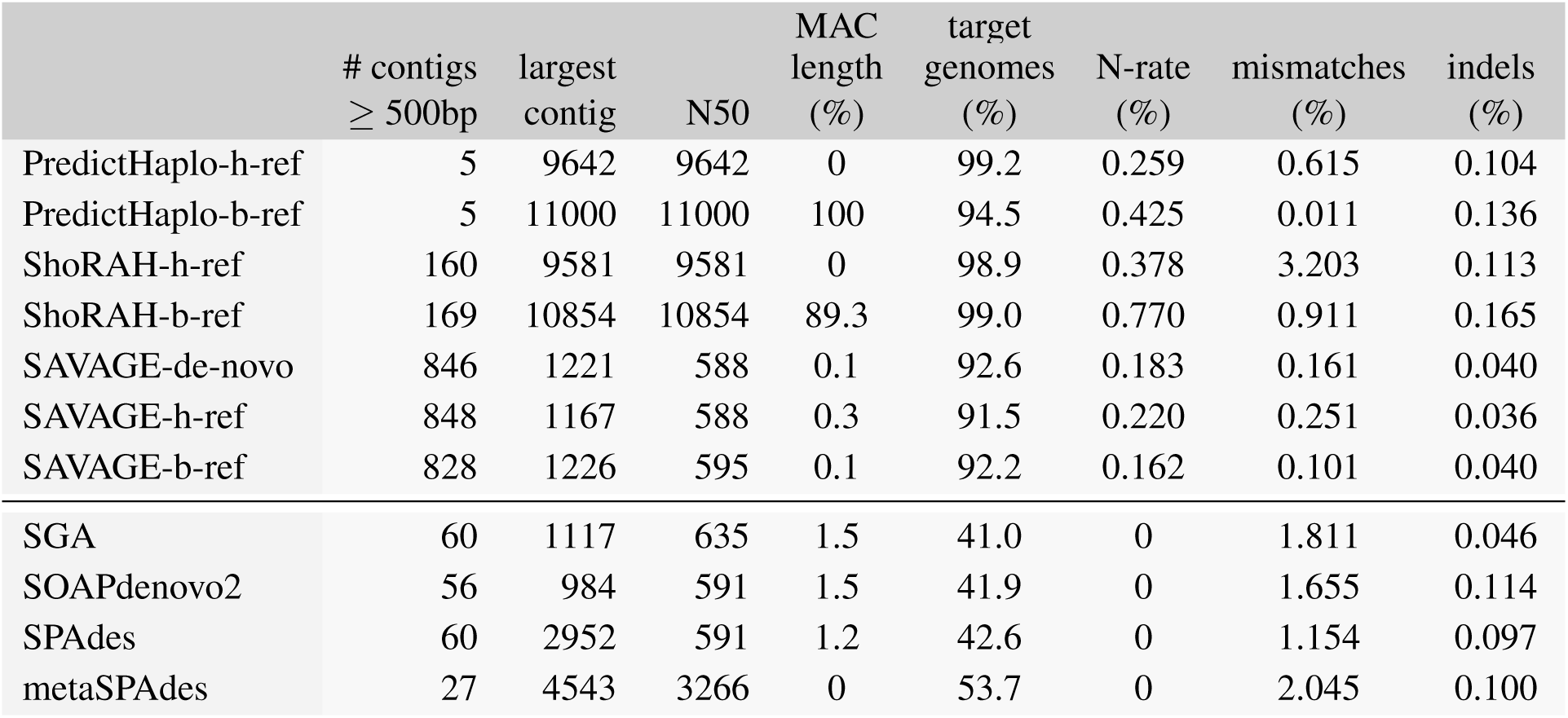
Assembly results per method on the HIV lab mix, a gold standard benchmark containing real sequencing data (20 000x coverage). For reference guided methods we present results using an established, high quality reference genome (h-ref) as well as an ad-hoc, bootstrap reference genome (b-ref). All assemblies were evaluated on the following criteria: number of contigs ≥ 500 bp, length of the largest contig, N50 statistic, MissAssembled Contigs (MAC) length relative to total contig length, percentage of the target genomes recovered, percentage of undetermined bases (N), and percentage of mismatches and indels compared to the ground truth.

| | # contigs<br>$\geq 500\text{bp}$ | largest<br>contig | N50 | MAC<br>length<br>(%) | target<br>genomes<br>(%) | N-rate<br>(%) | mismatches<br>(%) | indels<br>(%) |
| --- | --- | --- | --- | --- | --- | --- | --- | --- |
| PredictHaplo-h-ref | 5 | 9642 | 9642 | 0 | 99.2 | 0.259 | 0.615 | 0.104 |
| PredictHaplo-b-ref | 5 | 11000 | 11000 | 100 | 94.5 | 0.425 | 0.011 | 0.136 |
| ShoRAH-h-ref | 160 | 9581 | 9581 | 0 | 98.9 | 0.378 | 3.203 | 0.113 |
| ShoRAH-b-ref | 169 | 10854 | 10854 | 89.3 | 99.0 | 0.770 | 0.911 | 0.165 |
| SAVAGE-de-novo | 846 | 1221 | 588 | 0.1 | 92.6 | 0.183 | 0.161 | 0.040 |
| SAVAGE-h-ref | 848 | 1167 | 588 | 0.3 | 91.5 | 0.220 | 0.251 | 0.036 |
| SAVAGE-b-ref | 828 | 1226 | 595 | 0.1 | 92.2 | 0.162 | 0.101 | 0.040 |
| SGA | 60 | 1117 | 635 | 1.5 | 41.0 | 0 | 1.811 | 0.046 |
| SOAPdenovo2 | 56 | 984 | 591 | 1.5 | 41.9 | 0 | 1.655 | 0.114 |
| SPAdes | 60 | 2952 | 591 | 1.2 | 42.6 | 0 | 1.154 | 0.097 |
| metaSPAdes | 27 | 4543 | 3266 | 0 | 53.7 | 0 | 2.045 | 0.100 |

Recently, the first specialized *de novo* assembler has appeared (Malhotra et al. 2016b). We ran this method, called MLEHaplo, on our benchmarking data sets. Unfortunately, it could only handle the 600x HIV mix; for all 20 000x benchmarks, MLEHaplo did not finish within a week and used more than 140GB of main memory per data set. On the 600x HIV mix, it performed very poorly, reconstructing only 10% of the target genomes at a mismatch rate of more than 2%.

### Dependence of reference based approaches on reference genome quality

Reference based quasispecies assembly tools proved to perform adequately when a high quality reference genome is available (Zagordi et al. 2011; Prabhakaran et al. 2014). We question whether reference based approaches could yield appropriate quasispecies assemblies if provided with a *de novo* assembled genome sequence obtained from the sample reads, rather than a high quality reference genome. To address this point, we compared state-of-the-art methods PredictHaplo (Prabhakaran et al. 2014) and ShoRAH (Zagordi et al. 2011) on our benchmarks (Table 1) in two settings: either with a high quality reference genome, or with a genome sequence obtained by running the VICUNA assembler (Yang et al. 2012) on the sample reads. We refer to the former as a **high quality reference genome**, denoted *h-ref*, and the latter as a **bootstrap reference genome**, denoted *b-ref*. The quality of the output assemblies, as evaluated with MetaQUAST, is described in Tables 2 and 3, as well as Supplemental Tables S1 and S2.

For PredictHaplo and ShoRAH, the number of output sequences provides an estimate of the total number of strains in the quasispecies, since each output sequence represents a putative strain in the quasispecies. In Table 2, we see that on all benchmarks except the 15-strain ZIKV mix, the number of output sequences for PredictHaplo is very close to the true number of strains. For the 3-strain ZIKV mix, both the high quality reference genome and the bootstrap reference genome lead to a perfect assembly of 3 sequences without any mismatches and less than 0.042% indels (Supplemental Table S2). But considering the remaining (more challenging) data sets, we see that using a bootstrap reference genome causes a serious loss in the fraction of target genomes recovered by PredictHaplo (compared to using a high quality reference). On the 600x HIV mix and the lab mix, using the bootstrap reference even results in 100% of the sequences being misassembled (Supplemental Table S1). Only for the 15-strain ZIKV mix the difference between the h-ref and b-ref approaches is small: both recover only 53% of the target genomes (8 out of 15 strains – see Table 2).

For ShoRAH, we observe that for all data sets the number of output sequences is one or two orders of magnitude larger than the true number of strains. In addition, the mismatch rate is high compared to other methods, varying between 2.4% and 4.4% on the simulated 20 000x benchmarks. Unfortunately, we can only compare the bootstrap reference and high quality reference approaches on the HIV data, because ShoRAH-h-ref crashed repeatedly on the HCV and ZIKV benchmarks. Remarkably, the bootstrap reference approach increases target genome coverage from 39.4% to 93.8% on the 5-strain HIV mix (Table 2). However, in both the 20 000x HIV mix and the 600x HIV mix we see that the bootstrap reference also results in a small fraction of the total sequence length being misassembled (7.0% and 1.6%, respectively). This effect becomes much more apparent on the lab mix, with 89.3% of the total sequence length being misassembled. This shows that, similar to PredictHaplo, the quality of the ShoRAH assembly is highly dependent on that of the reference genome sequence.

Both tools, especially PredictHaplo, seem valuable when the reference genome is closely related to sample strains, but inadequate to handle cases where a good reference genome is unavailable. Moreover, Figure 2 shows that both PredictHaplo and ShoRAH have trouble reconstructing low-frequency strains, recovering less than 17% of the low-frequency (<5%) target strains. These results emphasize the need for new assembly approaches that are independent of a reference genome.

### SAVAGE evaluation

For the sake of comparison, we ran SAVAGE on the same benchmarks as above (Table 1) in both *de novo* mode and reference mode, both with default parameters. The 20 000x coverage data sets were split into patches of 750x each, on which we applied SAVAGE *Stage a*. Subsequently, all *Stage a* contigs were put together into one big collection of contigs and used as input for *Stage b* (Supplemental Figure S1).

Table 2 presents the evaluation results on simulated benchmarks of the *Stage b* maximally extended contigs for each of the three modes: SAVAGE-h-ref with a high quality reference genome, SAVAGE-bref with the genome assembled by VICUNA, and SAVAGE-de-novo (without reference). Remember that all *de novo* assemblers, including SAVAGE, proceed by progressively assembling longer and longer contigs starting from the raw reads, until finally, each output contig may (partially) cover the target genomes. Hence, unlike for PredictHaplo and ShoRAH, the number of contigs cannot be interpreted directly as a number of strains.

With a reference, the results of SAVAGE-h-ref and SAVAGE-b-ref are very similar: the contigs cover more than 99% of the target genomes, with the largest contig length close to the genome size of the virus in question. The mismatch, indel, and N rates are globally better than those offered by PredictHaplo and ShoRAH: the indel and N rates are respectively one or two orders of magnitude lower. Above all, the contigs are free of misassemblies (MAC length is 0%). Strikingly, providing a high quality reference genome or a bootstrap genome makes little difference, and on some datasets SAVAGE with a bootstrap genome achieves better results for certain statistics (higher N50, larger target genome fraction, lower mismatch rate for the 15-strain ZIKV mix in Table 2). These observations also hold on the lab mix (Table 3), where SAVAGE-ref recovers 91.5–92.2% of the target genomes at a mismatch rate of 0.101–0.251% and very low indel rates.

On all benchmarks, SAVAGE-de-novo delivers an assembly that is qualitatively at least as good as the SAVAGE-h-ref and -b-ref assemblies. Figure 2 shows that, in terms of target genome recovered, SAVAGE-de-novo slightly but consistently outperforms SAVAGE-ref. More importantly, this figure shows that both SAVAGE-de-novo and SAVAGE-b-ref greatly outperform *all* other methods, especially on low-frequency strains (i.e., frequency < 10%).

To analyze the effect of read length on SAVAGE assembly performance, we also built a 5-strain HIV mix with the exact same properties as given in Table 1, but with shorter reads (2×150 bp). We evaluated the resulting maximally extended contigs for SAVAGE-de-novo, SAVAGE-h-ref, and SAVAGE-b-ref (Supplemental Table S3). Compared to the original 5-strain HIV mix, which has 2×250bp reads, SAVAGE produces a more fragmented assembly but still covers 90.6–98.4% of the target genomes with mismatch rates between 0 and 0.006%.

Overall, SAVAGE can process samples containing a mixture of multiple strains and recover most of the target genomes with a high level of sequence quality. It performs slightly better in *de novo* mode than with a reference sequence and also performs well on shorter sequencing reads. Moreover, compared to existing methods, our approach does not suffer from misassemblies. For SAVAGE-de-novo, the misassembled contig (MAC) length is 0% on all simulated datasets and 0.1% on the lab mix, which drastically outperforms all approaches that reach ≥ 90% genome coverage and operate *without a high quality reference*. Moreover, SAVAGE can take advantage of a bootstrap reference sequence built by a single genome assembler. Finally, SAVAGE offers contigs with improved mismatch and indel rates, especially on low-frequency strains.

### Runtime and memory usage

We evaluate algorithm efficiency on both the 600x and the 20 000x simulated HIV mix, as well as the lab mix. We report CPU time and maximum memory usage for all methods evaluated previously on each of these HIV data sets in Supplemental Table S4. In terms of CPU time, SAVAGE-b-ref was considerably faster than SAVAGE-de-novo, with 6.4 versus 19 minutes on the 600x HIV mix, 449 versus 5296 minutes on the 20 000x HIV mix, and 850 versus 7495 minutes on the *lab mix*. This was to be expected, since *de novo* overlap graph construction requires enumeration of all approximate suffix-prefix overlaps among the reads. In comparison, PredictHaplo was faster but of the same order of magnitude as SAVAGE-b-ref with 7, 223, and 158 minutes, respectively. ShoRAH was comparable to SAVAGE and PredictHaplo on the 600x HIV mix (12 min) but very slow on the 20 000x data (22256–32375 min). The de Bruijn graph-based assemblers (SOAPdenovo2, SPAdes, and metaSPAdes) were very fast on all data sets, with a CPU time of 0.15–2 minutes on the 600x HIV mix, 5–46 minutes on the 20 000x HIV mix, and 6–166 minutes on the lab mix. The generic assembler SGA was considerably slower, with 24, 164, and 300 minutes, respectively. Finally, with 54 minutes on the 600x data MLEHaplo was the slowest, which also points out why it could not finish the 20 000x benchmarks.

Peak memory usage varied between 0.04 GB (PredictHaplo) and 8.4 GB (SPAdes/metaSPAdes) for the 600x HIV mix, between 0.5 GB (SGA) and 10 GB (ShoRAH) for the 20 000x HIV mix, and between 0.7 GB (SGA) and 12 GB (ShoRAH) for the lab mix. Both SAVAGE-de-novo and SAVAGE-b-ref are on the lower end of this scale, with 0.6/1.3 GB for the 600x HIV mix, 0.9/1.7 GB for the 20 000x HIV mix, and 1.1/3.0 GB for the lab mix, respectively. A complete comparison of runtime and memory usage for all methods is presented in Supplemental Table S4.

### Effect of strain divergence and relative abundance

Assembling the sequences of several strains from a viral sample may turn out more difficult depending on both the level of strain divergence and on their relative abundance. After comparing SAVAGE to state-of-the-art methods, we investigated the ranges of divergence levels and of relative abundances that SAVAGE can properly handle, and examined the combined effect of these two parameters on the assembly quality. We used a series of 36 benchmark datasets simulated from two HIV-1 strains: a combination of six divergence levels (from .5 % until 10% of nucleotidic divergence) with six ratios of abundance (from 1:1 until 1:100). We ran SAVAGE-de-novo and SAVAGE-b-ref (i.e., with VICUNA assembled genome). All assemblies were evaluated with MetaQUAST, and Figure 3 reports the heatmaps of (A) the coverage fraction of the two genomes, (B) the mismatch rate, and (C) the relative error on the frequency estimates of each strain.

**Figure 3:**
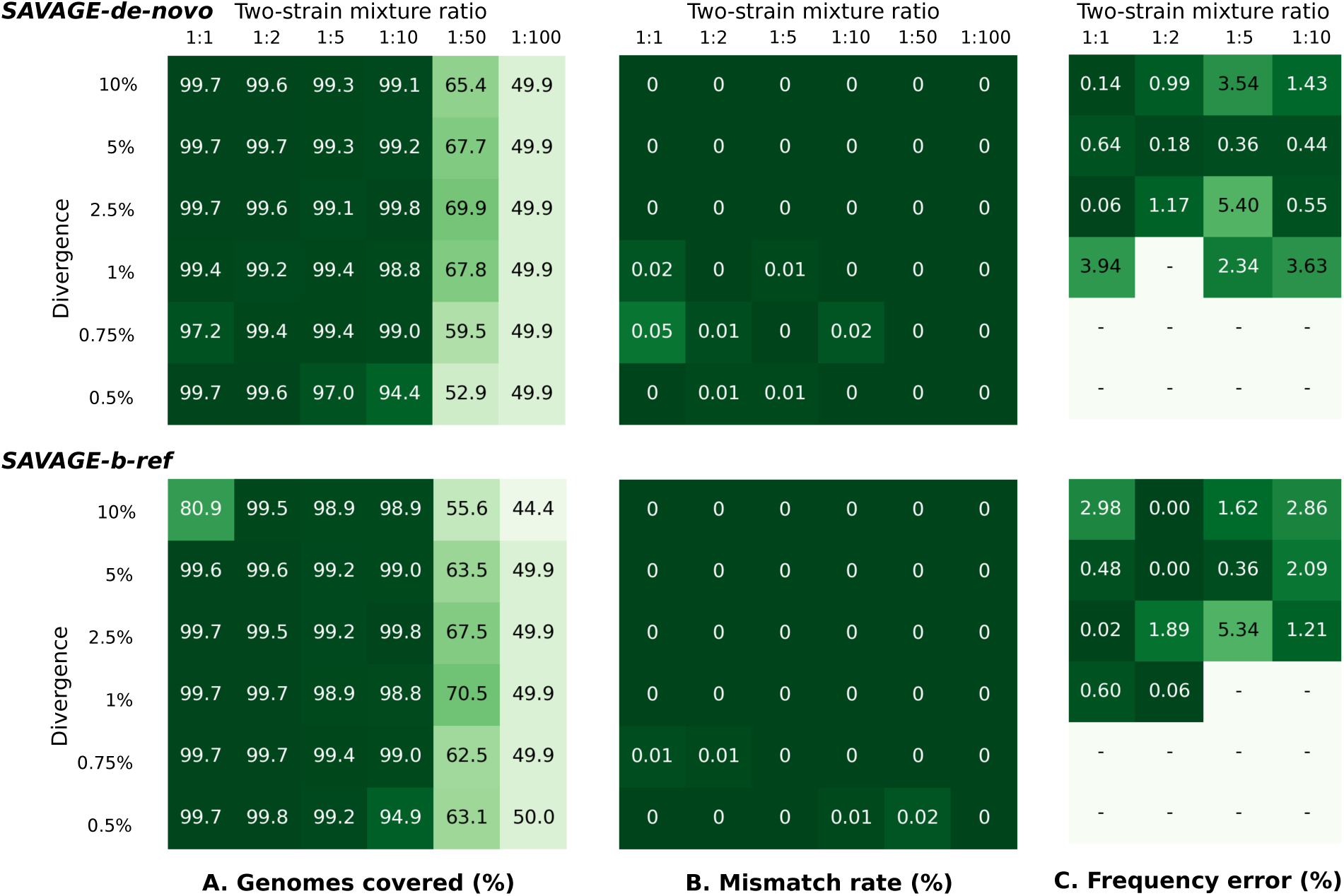
Performance of SAVAGE-de-novo and SAVAGE-b-ref, depending on pairwise distance and mixture ratio. **A.** Target genome fraction recovered (%) considering all maximally extended contigs ≥ 500 bp. **B.** Overall mismatch rate (%) considering all maximally extended contigs ≥ 500 bp. **C.** Relative error of estimated frequency for the minor strain (%). Frequency estimates were computed using Kallisto and only assemblies containing exactly two maximally extended contigs longer than 4000 bp were evaluated.

Comparing the two modes of SAVAGE, *de novo* or with a bootstrap reference, we observe similar results and a slight advantage to SAVAGE-de-novo in terms of genomes coverage. Altogether, SAVAGE obtains quasispecies assemblies of very low mismatch rates for all divergence levels and all relative abundance ratios, proving its ability to distinguish sequencing errors from true mutations. In general, the target genome coverage is very high for relative abundance ratios starting from 1:1 until 1:10, at all divergence levels. As the relative abundance of the minor strain decreases, it becomes more difficult to reconstruct the corresponding sequence. An extreme relative abundance of 1:100 hinders SAVAGE to reconstruct both strains: genome coverage values around 50% indicate that only one of the two strains has been assembled. We conclude that SAVAGE performs well in both modes (*de novo* and reference-guided) for relative abundances above 1:50 and a wide range of divergence levels.

#### Capacity to estimate the frequency of each strain

The problem of estimating relative frequencies of the contigs assembled for a viral quasispecies is very similar to quantifying the abundances of bacterial genomes from HTS data. Previous work (Bray et al. 2016) has shown that Kallisto can accurately tackle the latter problem, so we applied this method to our virus contigs as well (see Methods). For the 36 synthetic ‘divergence-vs-ratio’ benchmarks, we compared the estimated frequency of the minor strain in the sample with the real frequencies. The rightmost panel of Figure 3 shows the relative difference between the estimated frequency and the true frequency of the minor strain. This comparison was performed only when the strains were almost fully assembled (exactly two strains of length ≥ 4000 bp), hence abundance ratios of 1:50 and 1:100 were excluded. Of the remaining 24 datasets, 9–10 samples did not satisfy these criteria; the corresponding entries are marked ‘-’ in the heatmaps. Since there are only two strains in the sample, the absolute error is identical for both strains; however, the relative error will be much larger on the low-frequency strain. Hence, we evaluate performance on the most difficult task, namely estimating the frequency of the minor strain. In general, the relative estimation errors are very low: on average 1.65% for the SAVAGE-de-novo contigs and 1.39% for the SAVAGE-b-ref contigs, with an overall minimum of 0% (a perfect estimate) and a maximum of 5.34%.

### Zika virus sample

To test SAVAGE-de-novo on real conditions, we ran it on a sample taken from a rhesus macaque infected by an Asian lineage Zika virus (Dudley et al. 2016). The sequencing reads covered the full ZIKV reference genome used (NCBI sequence KU681081.3) at an average coverage of 30 000x. Using a similar procedure as for the real HIV data (lab mix), we split the reads into patches of approximately 750x each and proceeded with *Stage a* assembly on each patch (Supplemental Figure S1). Subsequently, we used the whole collection of *Stage a* contigs together as input for *Stage b*, which yielded 148 maximally extended contigs longer than 500 bp. A small fraction (4%) of these contigs could not be aligned to the reference genome, but instead matched four human BAC clones (accession AC117500.13, AC002565.1, AC079754.4, and AC015819.5) and one rhesus macaque BAC clone (accession AC190318.8) at *>* 90% sequence identity, indicating contamination, so we removed them from further consideration. The remaining 142 contigs contained 13 sequences longer than 1000 bp, the largest contig being 1874 bp long, and the N50 measure was 572 bp. The contigs covered the 10767 bp reference genome between positions 225–10767, the greatest divergence occurring between positions 1700 and 4200.

In *Stage c*, we allowed up to 1% divergence between contigs in the overlap graph, thus assembling representatives for groups of very closely related strains (see Methods). This resulted in 6 contigs of length at least 500 bp, now called master contigs. The largest sequence was 4155 bp long and the N50 measure was equal to 2065. Aligning the contigs to the reference genome reveals that the master contigs together form two master strains: their sequences differed only by a one nucleotide deletion at position 4103 followed by a SNP at position 4106 (see Supplemental Figure S2). Our frequency estimation procedure predicted the haplotype harboring the deletion to be the minor haplotype with a frequency of 8.6%, compared to 91.4% for the major haplotype. We hope that in the future, novel external data obtained by different means will become available for this sample, allowing an in-depth validation of our two-strain quasispecies assembly.

### Hepatitis C virus sample

Analogous to the ZIKV analysis above, we applied SAVAGE to a hepatitis C patient sample presented in (Töpfer et al. 2014). This sample covers the NS5B region (positions 7602–9374), a gene encoding for the RNA-dependent RNA polymerase, which is essential for viral replication. We found 839 contigs in *Stage b*, with an N50 measure of 533 bp and the largest contig 839 bp long. Aligning the contigs to the HCV reference genome (NCBI sequence NC_004102.1) reveals that the 9646 bp genome was covered between positions 6128–9304, with a relatively constant amount of variation across the whole region. We observed no contigs resulting from sample contamination (all contigs could be aligned to the reference sequence).

By allowing up to 1% divergence between contigs in the overlap graph in *Stage c*, we continued the assembly. This led to 80 master contigs of length at least 500 bp, of which 5 were longer that 1000 bp. The N50 measure was 535 and the largest sequence counted 1433 bp. Aligning the master contigs to the reference genome shows that one of the master contigs contains a large deletion of 444 bp. This particular sequence could not be aligned across the deletion; instead we found two clipped alignments for the contig, one for the first 781 bases and one for the last 319 bases. Combining these two alignments, the contig covers positions 7723–9267 the reference genome (nearly the entire NS5B gene), apart from a gap of 444 bases starting at position 8504. The largest master contig spans almost the same region (positions 7923–9356), but it does not show any deletions compared to the reference genome. We conclude that there is a 444 bp deletion in the NS5B gene of only a part of the strains in the sample, in agreement with results from an earlier study (Töpfer et al. 2014).

Compared to the previous sample (ZIKV), the current sample shows much more variation in both contigs and master strains. A likely explanation for this is the large difference in numbers of days of infection between the samples: 4 days for ZIKV versus 135 for HCV. To get an estimate on the number of master strains in the HCV sample, we built a conflict graph based on the alignments of the master contigs to the reference genome. An edge in this graph reflects that two contigs disagree on at least one position of the reference genome, hence any clique corresponds to a set of sequences all belonging to different strains. The largest clique in this graph was of size 16, suggesting the existence of at least 16 different strains in the HCV sample.

## Discussion

Recent outbreaks of viral diseases, such as the Ebola or the Zika virus, have pointed out a pressing need for methods to assess the genetic diversity of viral infections in a flexible manner, without strongly depending on the quality of available reference genomes. Here, we have presented SAVAGE, the first method for *de novo assembly of viral quasispecies* based on overlap graphs.

Viral genomes are characterized by high mutation and recombination rates. They are therefore often extreme in terms of both ploidy and the low relative abundance of single haplotypes. In our experiments, existing genome assemblers that do not depend on reference genomes were unable to reconstruct a viral quasispecies completely, where the (often resistance-inducing) low-frequency strains could not be captured sufficiently well. This has pointed out that only more specialized assemblers that can operate without depending on a reference genome have the power to overcome the current limitations.

We have shown that SAVAGE has this power and thus provides answers to such currently pressing issues. SAVAGE has performed very favorable—if not crucially advantageous—in comparison to a large collection of state-of-the-art de novo assemblers and specialized (but reference-dependent) viral quasispecies assemblers. Thereby, it proved particularly beneficial when being compared to reference-free approaches in terms of reconstructing strains of low frequency, which had been one of the essential goals of this study. Comparisons with existing reference-guided approaches pointed out that those yield contigs that are affected by more sequencing errors in general. Moreover, they tend to become confused by reference genomes of suboptimal quality, while SAVAGE behaves in a robust manner and can also make favorable use of such suboptimal bootstrap (ad-hoc) reference genomes. Last but not least, our method significantly outperforms the only available *de novo* viral quasispecies assembler (MLEHaplo) in terms of assembly quality, runtime, and memory usage. In an overall account, SAVAGE has proven to bridge a significant gap in the spectrum of viral quasispecies assembly approaches.

We believe that the central methodical reason for the benefits of our approach is the use of overlap graphs as the underlying assembly paradigm. While assembling genomes of low ploidy usually works favorably based on de Bruijn graphs, we have pointed out that using reads at their full length is key in assembling viral quasispecies, where distinguishing between low-frequency mutations and sequencing errors is imperative. The key insight is that (genetically linked) true mutations co-occur among different reads. Examining the full read span decisively enhances the detection of patterns of co-occurrence. Beyond enabling the detection of low-frequency strains, this also allows correction of sequencing errors in novel ways. We have pointed this out by making integrative use of sound statistical sequence models in combination with an iterative algorithmic scheme, which extends reads into contigs of increasing length and extremely low error content.

Key to reference free construction of overlap graphs has been the use of FM-index based techniques, which has been novel in the context of the analysis of viral data. Moreover, we have demonstrated that overlap graphs also seem to be the approach of choice when aiming to make use of ad-hoc consensus reference genomes, such as provided by specialized tools that construct a single consensus sequence from patient sample read data. Often, the resulting consensus sequence is of worse quality than a well-curated reference sequence. This can substantially disturb approaches that rely on the underlying reference as a sequence template (e.g. PredictHaplo, ShoRAH). Overlap graphs constructed by making use of reference sequence coordinates provide a robust alternative, since they use the reference sequence only as a coordinate system for the determination of overlaps.

A few more things are noteworthy. First, the bootstrap reference approach SAVAGE-b-ref has proven to outperform reference guided approaches in terms of the error rates of the contigs, even when they make use of high-quality reference sequence, which further underlines the general use of overlap graphs. Second, the target genome coverage of our full *de novo* approach SAVAGE-de-novo exceeded that of the high-quality reference-guided approaches, which points out its ability to distinguish sequencing errors from true mutations. Finally, SAVAGE-b-ref also depends on the quality of the reference sequence: the target genome coverage is 13.6 points lower compared to SAVAGE-h-ref on the 10-strain HCV mix. This, of course, had to be expected: if reference coordinates are too mistaken, overlaps cannot be detected. This last point underscores that a full de novo approach can come with decisive extra advanatages.

Of course, there is still room for improvements. While substantially faster and more space efficient than previous overlap graph based viral quasispecies assembly algorithms, SAVAGE has been particularly tailored towards dividing deep coverage datasets into chunks of 500 to 1000x, and merging the contigs of the chunks in subsequent steps, because this reflects its statistical calibration. While this works well, it sets certain limits on the frequency of strains it can recover — haplotypes of frequencies below 1% remain difficult to reconstruct. In future work, we will seek to lower these limits further by considering novel strategies for computing cliques in overlap graphs. On the algorithmic side, we will also explore alternative indexing techniques that allow for more relaxed definitions of overlaps and faster computation. Last but not least, incorporating long read data into SAVAGE may help to reconstruct full-length genomes.

## Methods

### Overlap graph construction

We first provide a brief definition of an overlap graph and then sketch how to construct such graphs from patient sample read data using *indexes* or *reference genomes* as two options.

#### Overlap graphs

For a collection 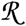 of sequencing reads (*Stage a*) or contigs (*Stages b,c*), both of which are sequences over the alphabet of nucleotides {*A, C, G, T, N*} (which includes *N* as a common placeholder for unknown nucleotides), the *overlap graph G* = (*V, E*) is a directed graph, where vertices *v* ∈ *V* correspond to reads/contigs 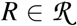 and directed edges connect reads/contigs 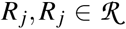 whenever a suffix of *R*_*i*_ of sufficient length matches a prefix of *R*_*j*_ and QS(*R*_*i*_, *R*_*j*_) ≥ δ where 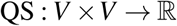 is a quality score that has to exceed a certain threshold δ. For *Stages a, b* we make use of the statistical model presented in (Töpfer et al. 2014), where QS(*R*_*i*_, *R*_*j*_) ≥ δ reflects that the overlapping parts of reads *R*_*i*_ and *R*_*j*_ present a locally identical haplotypic sequence. Note that the statistical model includes a refined analysis of the (Phredscaled) error profiles that underlie *R*_*i*_ and *R*_*j*_ so as to reflect that sequencing is an erroneous process and hence to assess the identity of their overlapping parts on a sound statistical basis.

In *Stage c*, QS(*R*_*i*_, *R*_*j*_) reflects the fact that the two contigs share only a limited amount of mismatches in their overlaps, meaning that they did likely emerge from identical master strain sequences.

#### Paired-end reads

SAVAGE was designed for short reads (typically Illumina reads); after merging self-overlapping pairs, the input in *Stage a* may contain paired-end reads and/or single-end reads. To make use of the pairing information, we add another edge restriction by allowing only the overlap cases shown in Figure 4. For overlaps involving a paired-end read, we require both read ends to have a sufficiently long overlap (at least half of the minimum overlap length for single-end reads) as well as a sufficiently high quality score.

**Figure 4:**
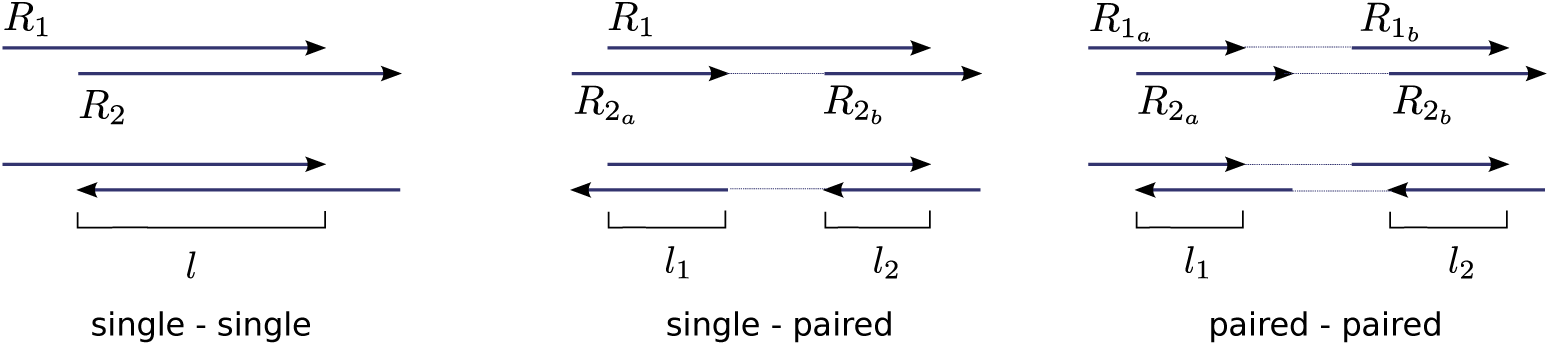
Edge criteria. For an overlap to become an edge in the overlap graph, it must satisfy three criteria. First, the overlap length *l* must be at least the minimal overlap length *L*. Second, the overlap quality score *QS*(*R*_1_*, R*_2_) must be at least the minimal score δ. For overlaps involving paired-end reads, we require both *l*_1_ ≥ *L* and *l*_2_ ≥ *L*, and, analogously, *QS*(*R*_1*a*_, *R*_2*a*_) ≥ δ and *QS*(*R*_1*b*_, *R*_2*b*_) ≥ δ. Finally, we only accept overlaps where the sequence orientations of a paired-end read agree: either both sequences in forward orientation, or both sequences in reverse orientation.

#### Construction

Construction of overlap graphs always proceeds in two steps. First, pairs of reads (*R*_*i*_, *R*_*j*_) are determined that have a sufficiently long and well-matching overlap. Subsequently, QS is evaluated on all pairs (*R*_*i*_, *R*_*j*_). For *Stages b and c*, where the input is sufficiently small, the first step is implemented by pairwise comparison of all contigs using BLAST (Altschul et al. 1990). The only difficulty is the first step in *Stage a*, where the input is very large (the original deep coverage data). This requires some sophistication; we explore two options:

1. *With a read index*: We determine all sufficiently long overlaps between sequencing reads using FM-index based techniques (Välimäki et al. 2012, SFO) such that overlaps contain at most 2% mismatches (accounting for up to 1% sequencing errors in each of the reads). This method, however, only works on single-end reads, so we first ignore the paired-end relations and consider each of the sequences as a single-end read. Then, after listing all pairwise overlaps with SFO, we reconsider the pairing information, outputting only overlaps that are supported by both read ends as described above.
2. *With a reference genome*: We align all reads against a reference genome; here we may use an ad-hoc consensus genome obtained by running an assembly tool on the sample reads. With all read alignments in hands, it is then computationally straightforward to determine all sufficiently long and sufficiently matching overlaps pairs.

#### Read orientations

When merging multiple reads into one consensus sequence, it is important that the reads agree on their respective orientations. Therefore, we apply a read orientation routine that assigns a label (+/−) to every read, indicating the orientation in which its sequence should be considered. This routine starts by setting the orientation of a node of minimal in-degree to +, then recursively labels all out-neighbors as defined by the corresponding edges (Figure 5, panel A). When there is no perfect labeling possible, meaning that there are conflicts among the read orientations due to inversions, we heuristically search for an orientation that leads to a minimal amount of conflicts among the reads.

**Figure 5:**
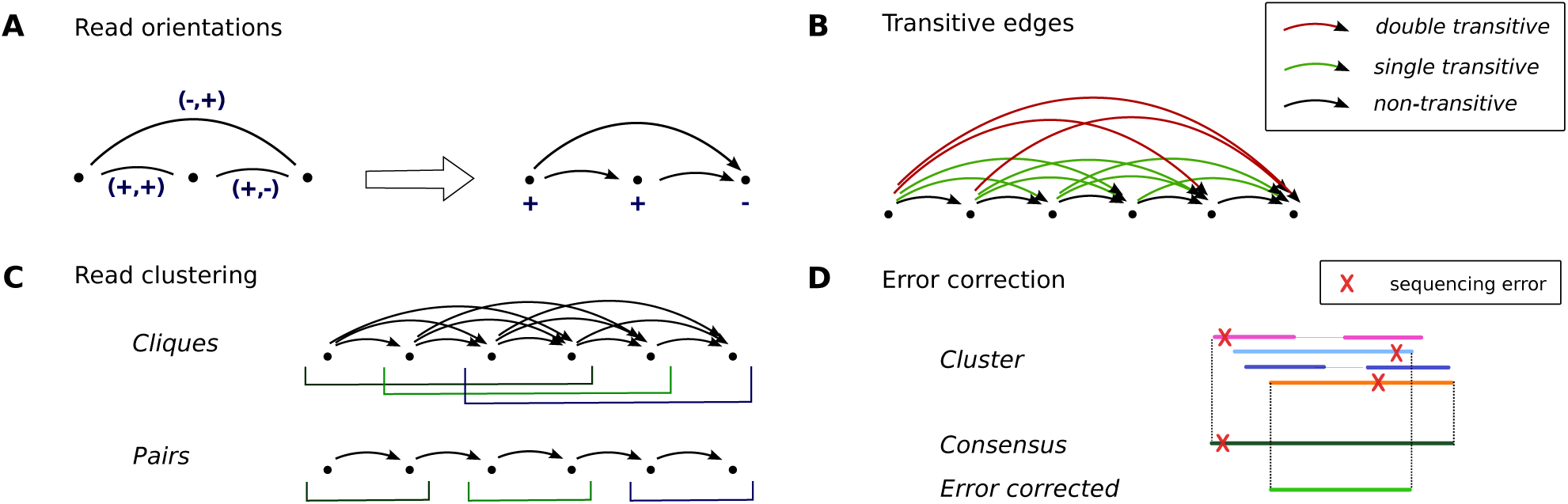
Algorithmic details. **A.** Read orientations: Given an edge *u* → *v* with orientations (−, +). Then if *u* is labelled +, the induced label for *v* is −, while if *u* is labelled − the induced label for *v* is +. This procedure leads to a vertex labeling in *O*(*V*) time. **B.** Transitive edges: An edge *u* → *w* is called *single transitive* (resp. *double transitive*), shown in green (resp. red), if there exists a vertex *v* and edges (resp. *transitive* edges) *u* → *v*, *v* → *w*. **C.** Read clustering by cliques (top) or by pairs (bottom). **D.** Error correction: when a consensus sequence is constructed from a cluster of reads, the extremities are removed.

### Overlap graph based assembly

In all stages, our algorithm proceeds as an iterative procedure where contigs grow with the iterations. The final contigs (in particular the output of *Stage b*, or, optionally, *Stage c*) can substantially exceed the length of the original reads. As our analyses demonstrate, these contigs present haplotype specific sequences with high accuracy.

#### Cliques and contigs

The main idea of our algorithm is to compute cliques in the overlap graph. A *clique* is a subset of the nodes such that each pair of nodes is linked by an edge. By definition of the edges, a clique groups reads that stem from identical haplotypes. Within a clique, reads/contigs share (possibly low-frequency) true mutations while sequencing errors are not shared by the majority of reads (Figure 1, Panel B). Hence, cliques can be used to clearly distinguish between true mutations and sequencing errors. This further allows us to correct these errors by transforming cliques into contigs that represent an error-corrected consensus sequence of the reads in the clique.

#### Transitive edge removal

The number of maximal cliques in an overlap graph grows exponentially with the number of nodes in the graph, that is here, with the read coverage of the dataset giving rise to the overlap graph. While our method relies on cliques for the purpose of error correction, the size of the cliques does not have to exceed a certain threshold for that goal.

A common approach to reduce the complexity of an overlap graph is to remove transitive edges (see e.g. (Simpson and Durbin 2012)). An edge *u* → *w* is called *transitive* if there exist a vertex *v* and edges *u* → *v*, *v* → *w*. We call an edge *u* → *w double transitive* if there exists a vertex *v* and *transitive* edges *u* → *v*, *v* → *w*, illustrated in Figure 5, panel B. Note that, by definition, any double transitive edge is also single transitive. We found that removing double transitive edges bounds the size of the cliques to 4, thus decisively limiting the number of maximal cliques and allowing efficient maximal clique enumeration, while still allowing for safely distinguishing errors from true mutations.

To find all double transitive edges, we first remove all non-transitive edges from the overlap graph to obtain the transitive graph *G*′. This can be done efficiently by computing the inner product of 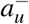 and 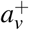 for all pairs (*u*, *v*) ∈ *V* × *V*, where 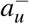 (resp. 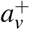) is the adjacency vector of outgoing (resp. incoming) edges of *u* (resp. *v*). Applying this procedure to *G* we obtain *G*′, and to find all double transitive edges we apply the same procedure to *G*′.

In the first iteration of *Stage a*, we remove all double transitive edges from the overlap graph. This reduces the number of contigs obtained in this iteration by an order of magnitude, leading to a decrease in CPU time and memory usage of even two orders of magnitude (Supplemental Table S5). In later iterations our algorithm no longer depends on clique formation because the reads (contigs) are assumed to be already of high quality. This allows us to remove not only double but also single transitive edges.

#### Read clustering

In the first iteration of *Stage a*, we cluster reads by enumerating maximal cliques in the overlap graph. After double transitive edge removal in an acyclic graph, the maximum clique size is 4, as illustrated in Figure 5, Panel B: a clique of size 5 will always use a double transitive edge. In practice, our overlap graphs are nearly acyclic and all cliques are of size at most 4. This implies that the total number of cliques is polynomial in the number of nodes, hence we can efficiently enumerate all maximal cliques; we use the degeneracy algorithm presented in (Eppstein et al. 2010) to do so. For the error correction algorithm to function optimally, we solely consider cliques of size 4 in this iteration.

In later iterations, after removing all single transitive edges, we merge pairs of contigs into new (extended) contigs. This does not require clique enumeration of any kind. See Figure 5, panel C for an illustration of the two read clustering techniques. In case of conserved regions among multiple strains, there can be branches in the overlap graph. In such situations it is often impossible to connect the variants left and right of the conserved region, hence we do not merge any pair of contigs connected by a branching edge (Supplemental Figure S3).

#### Contig formation and error correction

As outlined above, we transform all reads/contigs within a cluster (a clique or a pair of contigs) into a consensus sequence. It is important to determine the consensus very carefully, because the original sequencing reads may contain up to 1% sequencing errors. Every consensus base is determined by a position-wise weighted majority vote, where the weights correspond to the respective base quality scores, as described in (Töpfer et al. 2014) and Supplemental Methods. This procedure was designed to correct for all putative sequencing errors showing among members of a clique, which is especially relevant in the first iteration of *stage a* (the error correction step). In this specific iteration, therefore, we require cliques of size at least 4; it is then highly unlikely that all the reads in a clique will agree on a sequencing error. We remove the extremities of the resulting contig where the support of the clique is less than 4 (Figure 5, panel D). Reads that are not contained in any size 4 clique are discarded after this iteration.

#### Graph updating

The newly constructed contigs become the nodes of the updated overlap graph and we need to determine the edges between those nodes. In other words, we need to find all pairs of contigs satisfying our overlap criteria. In *Stage a*, we examine all pairs of contigs that share an original read. This approach is very efficient, but risks ignoring overlaps of contigs that do not share an original read. In *Stages b* and *c* the graph is sparse enough, such that we can update the edges by considering all induced overlaps. This means that for every edge *u* → *v* in the graph before updating, we consider every overlap *u*′ → *v*′ for all *u*′ ∈ *S*_*u*_, *v*′ ∈ *S*_*v*_, where *S*_*u*_, *S*_*v*_ are the sets of all newly constructed contigs containing *u*, *v*, respectively. In addition, we also reconsider all overlaps that were not included as an edge in the graph before updating due to an insufficient overlap quality score.

#### Iteration

The key idea of the SAVAGE assembly algorithm now is to repeatedly apply this twofold procedure of clique enumeration (*Stage a*) or merging pairs (*Stages b and c*) and contig formation. Thereby, all contigs of iteration *i* ≥ 1 become nodes in the overlap graph of iteration *i* + 1, which results in an overlap graph to be processed in iteration *i* + 1. We repeat this procedure until there are no more edges in the overlap graph. Key to success is that contigs are constantly growing along the iterations, and, upon convergence, greatly exceed the length of the original reads. An example of the progression of contig lengths during the three stages of the algorithm is given in Supplemental Table S6.

### Parameter settings

There are three parameters to be set, namely, the overlap score threshold δ, the mismatch rate *mr* allowed in the overlaps, and the minimal overlap length *L*. To analyze the behaviour of the overlap score function, we simulated 2×250 bp Illumina MiSeq reads from different genomes, diverging between 1% and 10%. We computed all overlaps among those reads and classified them by the number of true mutations in the overlap (not counting mismatches that are due to sequencing errors). This resulted in distributions *P*_*i*_, *i* ≥ 0, representing the overlap scores found in case of *i* true mutations (Supplemental Figure S4), from which we concluded that δ = 0.97 is the optimal choice. To be more conservative, this threshold can be raised, but this comes at the cost of a decrease in the target genome coverage.

The mismatch rate parameter allows overlaps having a sufficiently high overlap score to become edges in the overlap graph if the mismatch rate is sufficiently low. By default, this parameter is set to 0, meaning that we only rely on the overlap score for constructing the overlap graph. When assembling master strains, however, the allowed mismatch rate was set to 0.01, so that strains diverging by less than 1% were merged into a consensus sequence.

Finally, the setting of the minimal overlap length parameter depends on the average read coverage and sequencing depth. Increasing the minimal overlap length results in a faster algorithm and lower error rate, because the overlap graph will be very much restricted. But this achievement comes with a potential loss of low-frequency strains, since the corresponding reads may not have sufficiently long overlaps. In general, we found a minimal overlap length of 50–70% of the total read length to work well. The exact command lines and parameter settings used for all experiments can be found in Supplemental Methods.

### Frequency estimation

We apply Kallisto (Bray et al. 2016) to estimate relative frequencies of the contigs assembled for a viral quasispecies. Kallisto was designed for quantifying the abundances of bacterial genomes from HTS data, which is similar in spirit to estimating frequencies for viral quasispecies assembly. The Kallisto algorithm takes as input the original sequencing reads along with the contigs, and returns for every contig a so called TPM (Transcripts Per Million). This number estimates the amount of sequencing reads corresponding to this contig for every one million reads considered, and it is independent of the contig length. We translate these counts to relative frequencies by dividing each TPM by the sum of TPMs of all contigs evaluated. For the heatmaps in Figure 3, panel C, we only evaluated the two contigs of at least 4000 bp.

### Other methods used for evaluation

For benchmarking, we compared SAVAGE against the state-of-the-art approaches ShoRAH (Zagordi et al. 2011) and PredictHaplo (Prabhakaran et al. 2014). Both methods were run with default parameter settings, after aligning the reads to the reference genome using BWA-MEM (Li 2013). The *de novo* assembler MLEHaplo (Malhotra et al. 2016b) required the reads to be error corrected first, for which we used MultiRes (Malhotra et al. 2016a) with default settings (recommended by the authors). Unfortunately, we could not compare against VGA (Mangul et al. 2014) and HaploClique (Töpfer et al. 2014) because these software packages were no longer maintained.

### Data simulations

To evaluate performance of SAVAGE, we designed several simulated data sets. We used the software SimSeq (https://github.com/jstjohn/SimSeq) to simulate Illumina MiSeq reads from the genome of interest. In order to obtain reads similar to the real 5-virus-mix data, we simulated 2×250 bp paired-end reads, with a fragment size of 450 bp and the MiSeq error profile provided with the software. In addition, we also simulated a 5-strain HIV mix with shorter reads (2×150 bp). The genomes used for each data set are listed in the Supplemental Methods.

### Read trimming and merging

Before running any of the methods, the raw Illumina reads were trimmed using CutAdapt (Martin 2011). Next, we applied PEAR (Zhang et al. 2014) for merging self-overlapping read pairs. This resulted in a final read set containing both single-end and paired-end reads, on which we ran SAVAGE. For the other methods (MLEHaplo, PredictHaplo, ShoRAH, and VICUNA) we used the trimmed reads without merging, since neither of these methods accepts a combination of single- and paired-end reads. In addition, MLEHaplo required an error correction step on the input reads which was performed using MultiRes (Malhotra et al. 2016a).

### MetaQUAST evaluation

We use MetaQUAST (Mikheenko et al. 2016) for quality evaluation of the assembled contigs, which evaluates the contigs against each of the true viral genomes. By default, MetaQUAST uses the option `‐-ambiguity-usage all`, which means that all possible alignments of a contig are taken into account. However, the genomes in a viral quasispecies can be so similar that a contig may align to multiple strains, even though it only matches one haplotype. Therefore, we manually changed this option to `‐-ambiguity-usage one`, such that for every contig only the best alignment is used. Contigs shorter than 500 bp were ignored during evaluation.

### Data and software availability

A C++ implementation of SAVAGE is available for public use at https://bitbucket.org/jbaaijens/savage. The lab mix used for experiments can be downloaded from https://github.com/cbg-ethz/5-virus-mix and both the ZIKV and HCV data are available in the NCBI Sequence Read Archive under experiment SRX1678783 (run SRR3332513) and experiment SRX396803 (run SRR1056035), respectively. All simulated data sets can be downloaded from https://bitbucket.org/jbaaijens/savage-benchmarks.

## Acknowledgements

AS acknowledges funding through Vidi grant 639.072.309, provided by the Netherlands Organisation for Scientific Research (NWO). ER, AZ are supported by ANR Colib’read (ANR-12-BS02-0008), the Institut de Biologie Computationnelle (ANR-11-BINF-0002), and France Génomique.

### Disclosure declaration

The authors declare that they have no conflict of interest.

